# Chiral inversion mutagenesis identifies geometrically constrained residues within self-associating low-complexity domains

**DOI:** 10.64898/2025.12.17.694949

**Authors:** Ryan L. Beckner, Christien Carter, Glen Liszczak

## Abstract

Many protein low-complexity domains (LCDs) reversibly self-associate to enable cellular function, yet fundamental questions remain regarding how polypeptide chemical and structural features beyond side chain identity contribute to LCD-LCD interactions. For instance, the folds adopted by globular proteins emerge from constraints enforced by homo-chirality of genetically-encoded polypeptides. However, it remains unclear to what extent similar geometric constraints apply to LCD self-association. Herein, we use protein total- and semi-synthesis to probe the contribution of C^α^ stereochemistry to LCD self-association with synthetic Chiral Inversion Mutagenesis (ChIM). By introducing targeted L-to-D amino acid inversions, ChIM identifies C^α^ stereocenters under geometric constraint without modification of side-chain functionalities. We apply ChIM to the LCDs of inner nuclear lamina protein Emerin and neurofilament light chain and find that chiral inversion produces strongly position-dependent effects upon LCD self-association. Our study describes essential structural features that enable LCD self-association and chemical strategies to interrogate LCD biochemistry.

**Significance Statement:** This study identifies a critical role for amino acid side chain chirality in low-complexity domain (LCD) self-association. Using synthetic protein chemistry, we introduced targeted L-to-D amino acid inversions in LCDs without altering side chain functionalities. Chiral inversion mutagenesis identified geometrically constrained residues in the LCDs of Emerin and neurofilament light chain that localize to self-association hotspots within these sequences. Our work demonstrates that, like globular proteins, LCDs exploit polypeptide homochirality for biochemical function. The position-dependent effects of chiral inversion indicate the formation of labile structural elements that mediate oligomerization. By employing synthetic protein chemistry to probe LCD biochemistry with high positional resolution, this work provides a powerful approach to dissect chemical principles and polypeptide structural features that govern LCD-LCD interactions.

## Introduction

Protein low-complexity domains (LCDs) differ from traditional folded domains in that they are enriched in a small subset of amino acids and do not adopt stable three-dimensional structures. Due to their high solvent accessibility, LCDs are often hubs for protein-protein interactions and frequently serve as sites for regulatory post-translational modifications (1, 2). Additionally, despite low sequence complexity, oligomerizing LCDs drive diverse phenomena including intermediate filament (IF) assembly, the selective permeability of the nuclear pore barrier, and nucleic acid organization (3–6).

Although pathological LCD aggregation is known to produce both familial and sporadic neurodegenerative disease, the physiological mechanisms by which LCDs oligomerize into higher-order structures remain enigmatic. Composition, patterning, and valency of side chain functional groups are known to produce LCD self-association through the summation of several physiochemical interactions (7). However, recent evidence suggests that sequence-encoded and polypeptide backbone-mediated features robustly contribute to LCD self-association in addition to “bulk” physiochemical composition. For example, solid-state NMR (ssNMR) studies indicate that interacting disordered “head” domains of neurofilament light chain (NEFL) and desmin (DES) IFs become enriched in labile β-sheet structures templated by supramolecular filament assembly (8). In addition, independent studies have demonstrated that a transiently-structured region of RNA-binding protein TDP-43 contributes to self-association (9–11). Consistent with this principle, Eisenberg and co-workers have described low-complexity aromatic-rich kinked segments (LARKS) that adopt a kinked β-sheet architecture to enable reversible, sequence-encoded LCD self-association. These findings reinforce the idea that LCD oligomerization can arise via structurally privileged sequence motifs (12). Thus, interactions between discrete sequence-encoded elements can support physiological self-association of ostensibly disordered protein domains. Notably, recurrent pathogenic missense mutations in proline residues occur directly adjacent to these self-association hotspot sequences and enhance the strength of otherwise transient interactions to promote protein aggregation. LCD disease mutations of this class may therefore demarcate a local self-association-promoting element (9).

Interrogating sequence-encoded determinants of LCD self-association can be challenging due to the limited effect of conventional mutagenesis on LCD assembly (7, 13). In this context, methods adopted from synthetic protein chemistry have proven useful. In a previous study, our lab conducted scanning N^α^-methylation (“backbone”) mutagenesis of the TDP-43 LCD using semi-synthetic protein chemistry, revealing a short linear sequence wherein N^α^ methylation abrogates TDP-43 LCD self-association (9). Thus, synthetic mutagenesis of intrinsic polypeptide features beyond side chain identity can reveal unappreciated contributors to LCD self-association. To build on our previous study, we sought to assess how other fundamental polypeptide features contribute to LCD self-association. In folded proteins, polypeptide secondary and tertiary structural elements place geometric constraints on participating side chains (14). Indeed, stereochemical fidelity is a critical component of biomolecular structure-function paradigms, as well as a classic principle of chemical interaction specificity. However, it is unknown whether similar geometric rules apply to self-associating LCDs to achieve interaction specificity. We hypothesized that inverting residue C^α^ stereochemistry through synthetic L-to-D enantiomer substitution could be used to introduce a geometric perturbation without altering the “bulk” composition of side-chain functional groups (15, 16). This strategy, henceforth referred to as Chiral Inversion Mutagenesis (ChIM), can be applied to systematically scan LCD sequences for amino acid positions that are under geometric constraint during self-association. In this study, we apply ChIM to resolve chiral determinants of Emerin (EMD) LCD and NEFL head domain self-association.

## Results

### Delineation of a self-association hotspot in the EMD LCD

EMD is a small inner nuclear lamina protein that supports nuclear integrity in heart and skeletal muscle. EMD is comprised of an N-terminal LEM domain, an extended central LCD mediating self-association and lamin interaction, and a C-terminal transmembrane helix (Fig. 1A). Notably, recurrent mutations at Pro183(Thr/Arg/His) have been reported to cause X-linked Emery-Dreyfus muscular dystrophy (X-EDMD) through enhancing EMD self-association, mirroring recurrent LCD proline mutations found in familial neurodegenerative disease (17–19).

**Fig. 1.**
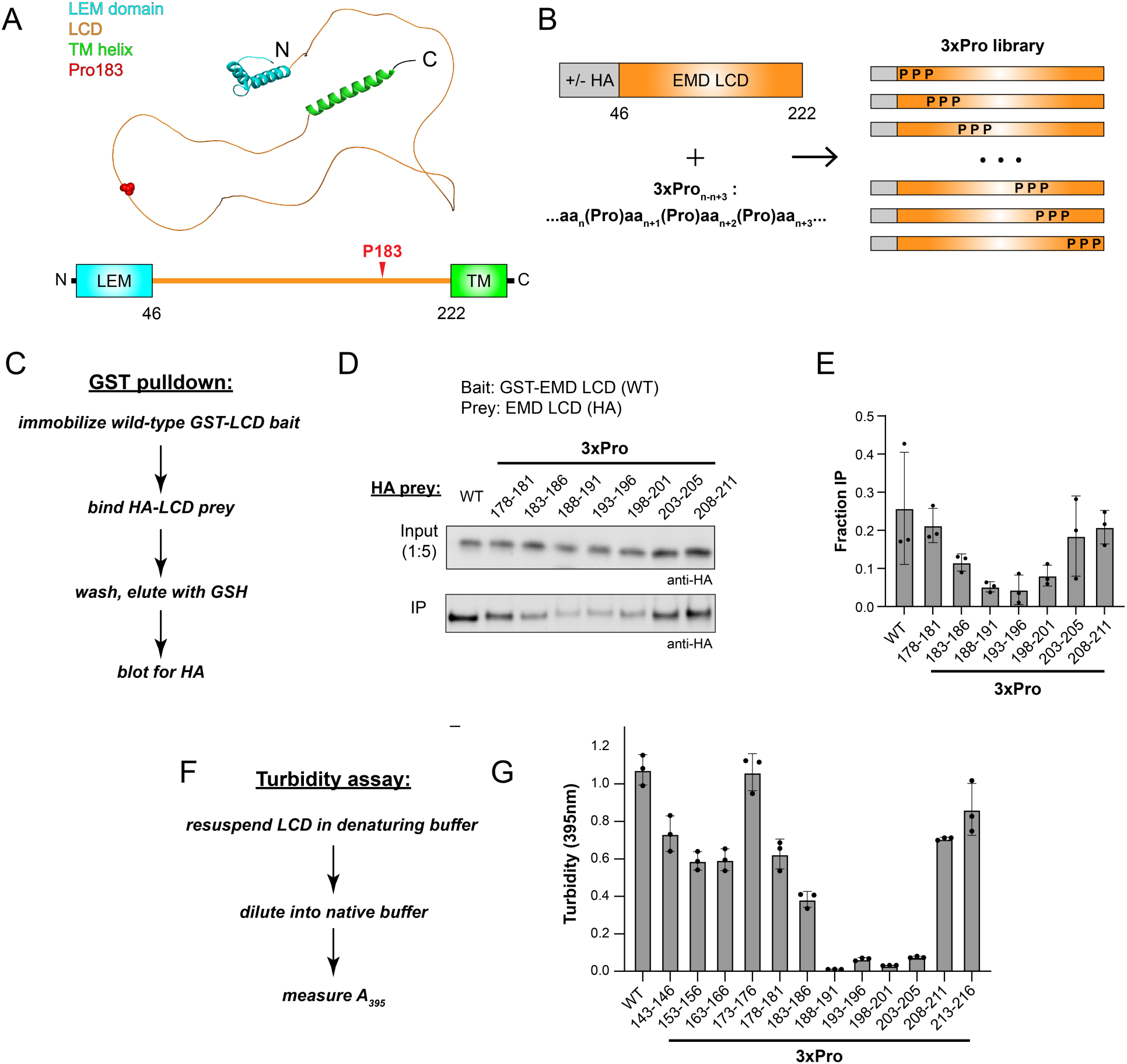
Delineation of a self-association hotspot in the EMD LCD. (A) AlphaFold-predicted structure of the full-length EMD protein and accompanying domain map. LEM domain (teal), LCD (orange), transmembrane helix (green), and aggregation-promoting disease mutation site Pro183 (red) are indicated. (B) Overview of the 3xPro NEFL library. (C) Workflow for the GST pulldown assay. (D) Representative Western blot from GST pulldown using HA-tagged 3xPro EMD LCD (50 nM) as prey. (E) Densitometry quantification of the GST pulldown assay shown in (D) (n=3). (F) Workflow for turbidity self-association assay (G) Turbidity-based EMD self-association assay quantification of untagged 3xPro EMD library (60 µM LCD, n=3).

To delineate the EMD LCD self-association hotspot, we devised a scanning 3x proline insertion (“3xPro”) mutagenesis campaign as an analogue of “backbone” mutagenesis (Fig. 1B). In this campaign, three proline residues were inserted in an alternating pattern with the native sequence. We first employed a recombinant protein-based GST-pulldown assay adapted from prior literature to probe 3xPro effects on EMD-EMD interactions (20). Briefly, wild-type glutathione S-transferase(GST)-EMD LCD fusion protein was immobilized on glutathione (GSH) resin, incubated with nanomolar concentrations of HA-tagged EMD LCD prey, washed with binding buffer, and eluted with GSH (Fig. 1C). Binding was assessed by Western blot against HA in eluent relative to input as quantified by densitometry. We found that 3xPro disrupts EMD-EMD interactions with strong position-dependence. The 3xPro between residues 188-201 (3xPro_188-191_, 3xPro_193-196_, 3xPro_198-201_) strongly abrogates pulldown by 5-fold, 5.9-fold, and 3.4-fold relative to wild-type, respectively. The 3xPro between residues 183-186 (3xPro_183-186_) modestly abrogates pulldown by 2.2-fold. The 3xPro between residues 143 and 183 or between residues 201 and 211 display similar pulldown to wild-type (Fig. 1D and E). Of note, the self-association hotspot identified using the 3xPro mutagenesis screen is directly C-terminal to the aggregation-promoting Pro183 disease mutation site.

Complementary turbidity assays were also performed on an extended panel of EMD 3xPro mutants. Transfer of LCDs from denaturing to native buffers through dilution or dialysis can produce turbidity due to LCD self-association, which can be quantified via absorbance at 395 nm (Fig. 1F) (9). Despite its marked hydrophilicity, the EMD LCD precipitates from solution at concentrations above 50 µM in native Tris buffer. We find that proline insertions within residues 188-206, including 3xPro_188-191_, 3xPro_193-196_, 3xPro_198-201,_ and 3xPro_203-206_, abrogate precipitation and reduce turbidity by 107.6-, 16.6-, 34.7-, and 14.3-fold, respectively (Fig. 1G). Proline insertion between residues 183-186 (3xPro_183-186_) modestly reduces turbidity by 2.8-fold relative to wild-type. Proline insertions N-terminal to residue 183 or C-terminal to residue 206 yield turbidity values within 2-fold of wild-type. In sum, turbidity assay data are in alignment with results from the GST pulldown wherein EMD residues 188-201 are necessary for the strong self-association of the EMD LCD. These data are consistent with previous truncation analyses proposing residues 170-220 as necessary for EMD homo-oligomerization (20).

### ChIM identifies enantioselective interaction hotspots in the self-associating region of EMD

To produce semi-synthetic EMD LCD variants for ChIM analysis, a recombinantly produced EMD-Piece 1 (EMD-P1) thioester fragment was ligated to a synthetic EMD-P2 peptide fragment bearing an N-terminal cysteine (Fig. 2). The EMD-P1 fragment (EMD residues 46-188) was expressed as a fusion protein with a C-terminal Gyrase A intein, purified via immobilized metal affinity chromatography (IMAC), and subjected to MesNa thiolysis followed by reverse-phase high performance liquid chromatography (RP-HPLC) purification. This EMD-P1 thioester fragment was produced with or without an N-terminal HA tag for GST-pulldown or turbidity assays, respectively. The EMD-P2 peptide fragment corresponding to residues 189-221 (S189C, D220E) was synthesized on Rink amide resin with microwave-assisted FMOC solid-phase peptide synthesis (SPPS) and purified with RP-HPLC. To systematically introduce C^α^ inversions through the EMD self-association core, a total of seven EMD-P2 synthetic fragments were prepared in which three D-amino acids (“3xD”) were incorporated at alternating positions in addition to all L- “wild-type” (Fig. 3A). Each EMD-P1 fragment was then ligated to each of the EMD-P2 fragments via trifluoroethanethiol (TFET) -assisted native chemical ligation and purified via RP-HPLC, yielding milligram-scale quantities of homogenous EMD LCD (residues 46-221) as characterized by SDS-PAGE and ESI-LC/MS (Fig. 3B, 3C S1, S2). We note that our EMD D-amino acid scan focused on the 189-221 segment, which was sufficient to cover both proline-sensitive and proline-insensitive regions. Additional sequence coverage would require multi-step ligations, which is particularly limiting for ChIM scanning-scale studies.

**Fig. 2.**
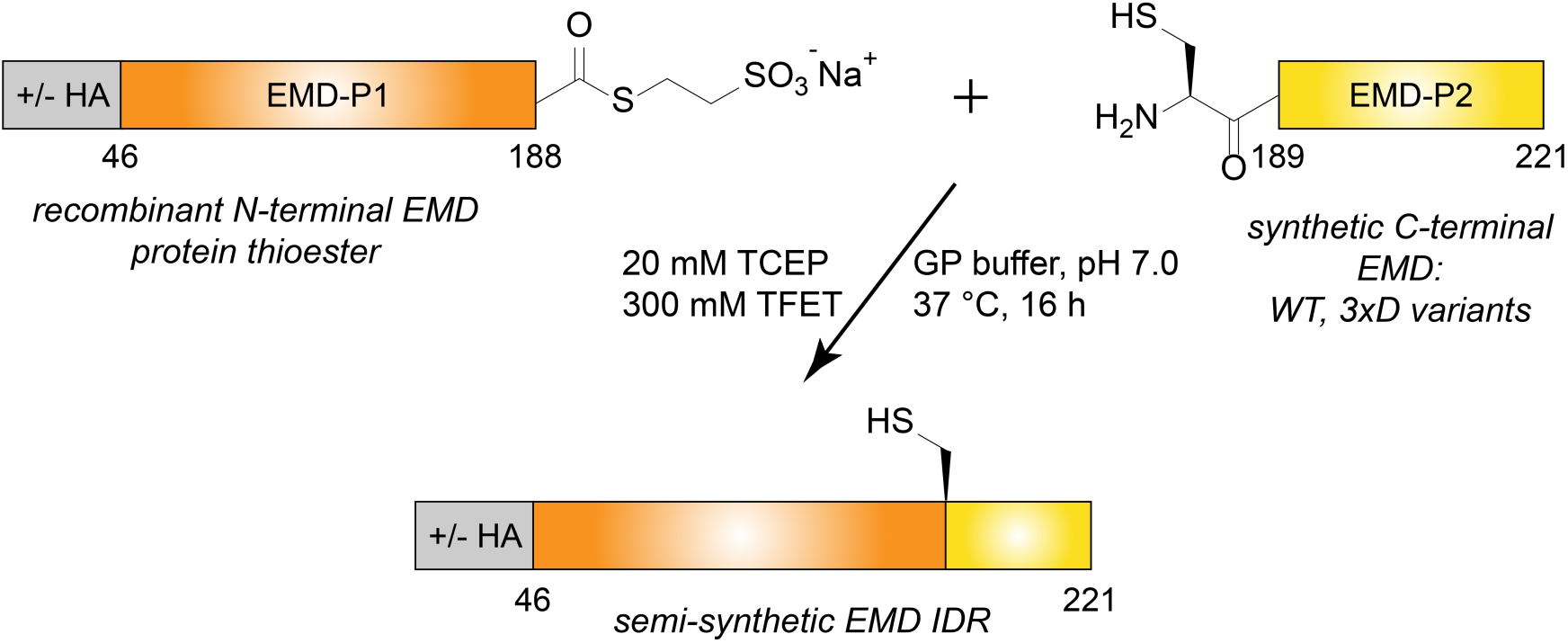
Semi-synthetic strategy to assemble the emerin LCD. The synthetic scheme used to assemble the EMD ChIM library. Note: HA = HA-tag; P1 = piece 1; P2 = piece 2.

**Fig. 3.**
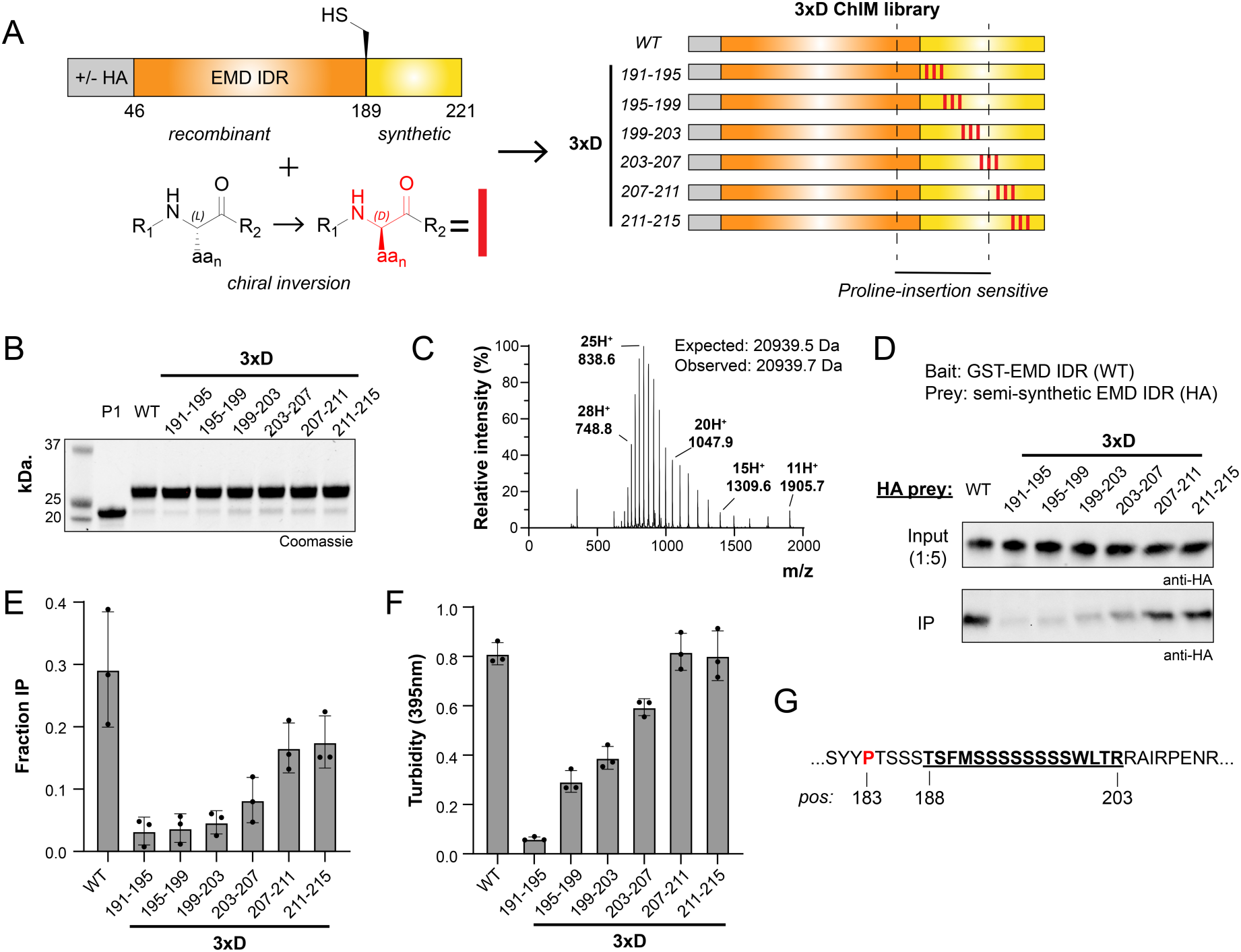
ChIM scanning identifies enantioselective interaction hotspots in the self-associating region of EMD. (A) Overview of the 3xD ChIM EMD library. The D-amino acid enantiomers were introduced into the synthetic portion of the LCD at the indicated positions (red rectangles). (B) SDS-PAGE characterization of HA-tagged 3xD ChIM EMD library. (C) Representative ESI-LC/MS spectrum of a semi-synthetic 3xD EMD LCD. (D) Representative Western blot from GST pulldown using HA-tagged 3xD EMD LCD (50 nM) as prey. (E) Densitometry quantification of GST pulldown assay shown in (D) (n=3). (F) Turbidity-based EMD self-association assay quantification of 3xD ChIM EMD library (50 µM LCD, n=3). (G) The EMD self-association hotspot sequence (underlined and bold) as defined in proline insertion mutagenesis and ChIM scanning. The disease mutation site Pro183 is shown in red. Note: ChIM scanning begins at residue 191.

To identify C^α^ stereocenters critical for EMD self-association, we conducted the GST-pulldown assay with our EMD 3xD ChIM library (Fig. 3D and E). We found that 3xD inversions within the EMD self-association hotspot identified in our 3xPro insertion scan (aa 188-201) strongly abrogate pulldown. These include 3xD_191-195_, 3xD_195-199_, and 3xD_199-203_, which respectively yield 8.8-, 7.9-, 6.2-fold pulldown reductions relative to wild-type. Construct 3xD_203-207_, which comprises 3xD inversions at the edge of the self-association hotspot, modestly abrogated pulldown with 3.6-fold relative reduction. Chiral inversion outside of the hotspot, constructs 3xD_207-211_ and 3xD_211-215_, retained pulldown values within 1.8- and 1.7-fold of wild-type.

Additionally, a series of untagged EMD LCDs with identical 3xD inversions was produced and subjected to the EMD self-association turbidity assay (Fig. 3F). Consistent with GST pulldown, residues within 191-203 are sensitive to chiral inversion with 13.3-, 2.8-, and 2.1-fold decreases in turbidity (3xD_191-195_, 3xD_195-199_, and 3xD_199-203_ respectively). Construct 3xD_203-207_ on the C-terminal edge of the hotspot displays a modest 1.4-fold decrease in turbidity while constructs 3xD_207-211_ and 3xD_211-215_ retain wild-type turbidity.

Despite conservation of number and sequence of side-chain functional groups, certain EMD 3xD diastereomers lack self-association. Therefore, privileged C^α^ stereocenters within the serine-rich EMD LCD are constrained during self-association (Fig. 3G). These data show that EMD LCD self-association exploits geometric features of the polypeptide backbone, consistent with formation or stabilization of a transiently structured element.

### ChIM scanning identifies self-association hotspots within the NEFL head domain

A prototypical class IV intermediate filament, NEFL is comprised of a disordered N-terminal head domain, a central coiled-coil rod domain, and a disordered C-terminal tail domain (Fig. 4A). Recurrent mutations at NEFL Pro8(Arg/Gln/Leu) and Pro22(Arg/Ser/Thr) in the head domain enhance self-association and cause familial Charcot-Marie-Tooth neuropathy (CMT) (21, 22). Characteristic of IFs, the rod domain of NEFL forms tetramers that assemble first laterally into unit-length filaments and subsequently longitudinally into mature filaments. Integrated IF superstructure, exemplified by the model class III IF vimentin (VIM), was recently revealed by cryo-EM as a ring of five octamer protofibrils with disordered head domains interacting in a condensed polymer within the filament lumen (Fig. 4B) (23). Extrapolating this model to NEFL, and considering prior ssNMR studies identifying labile β-structure within head domain polymers and assembled filaments, we applied ChIM analysis to map C^α^ constraint in the NEFL head domain.

**Fig. 4.**
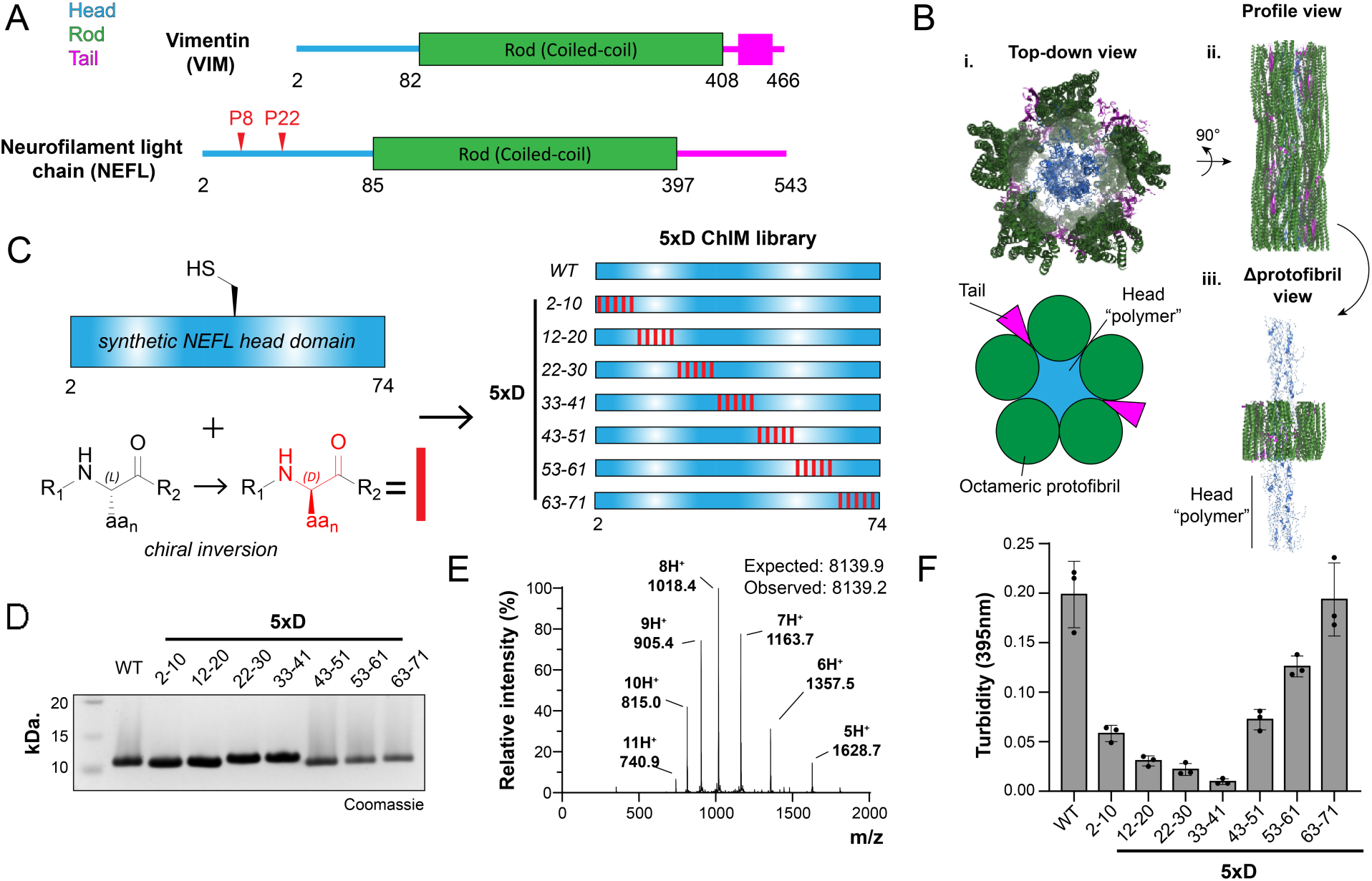
ChIM scanning identifies self-association hotspots within the NEFL head domain. (A) Domain maps of NEFL and VIM intermediate filaments. Head domains (blue), coiled-coil rod domains (green), tail domains (magenta) and aggregation-promoting NEFL proline mutation sites (red) are indicated. (B) Structure of VIM filament adapted from Ref. 23 (PDB:8rve). (i) Top view of VIM filament and corresponding cartoon representation, (ii) Profile view of the VIM filament. (iii) Profile view of VIM filament with internal “head domain polymer” exposed. (C) Overview of the 5xD ChIM NEFL library. The D-amino acid enantiomers were introduced into the synthetic LCD at the indicated positions (red rectangles). (D) SDS-PAGE characterization of the 5xD ChIM NEFL head domain library. (E) Representative ESI-LC/MS spectrum (left) of a synthetic 5xD NEFL head domain. (F) Turbidity-based NEFL head domain self-association assay quantification of 5xD ChIM NEFL head domain library (100 µM LCD, n=3).

Synthesis of the NEFL head domain was achieved by two-piece native chemical ligation (Fig. 5). A series of synthetic NEFL-P1 peptide fragments corresponding to residues 2-30 of NEFL were prepared with a C-terminal bis(2-sulfanylethyl)amide (SEA) moiety using modified FMOC SPPS (24, 25). Crude NEFL-P1 peptides were converted to MesNa thioesters and purified with RP-HPLC. Synthetic NEFL-P2 peptide fragments corresponding to NEFL residues 31-74 with an N-terminal cysteine (S31C, D72E) were synthesized with conventional FMOC SPPS and purified with RP-HPLC. D-amino acid inversions were introduced into either NEFL-P1 or NEFL-P2 fragments to perform a systematic ChIM scan through the entire NEFL head domain (Fig. 4C). A total of nine synthetic peptide fragments were prepared in which five D-amino acid inversions were incorporated at alternating C^α^ positions (“5xD”). A TFET-assisted native chemical ligation reaction was performed to ligate the NEFL-P1 and NEFL-P2 fragments, and the products were purified by RP-HPLC to yield a total of seven synthetic NEFL head domains. The purity of all synthetic NEFL head domain was determined by SDS-PAGE, ESI-LC/MS, and analytical HPLC (Fig. 4D, 4E, S3).

**Fig. 5.**
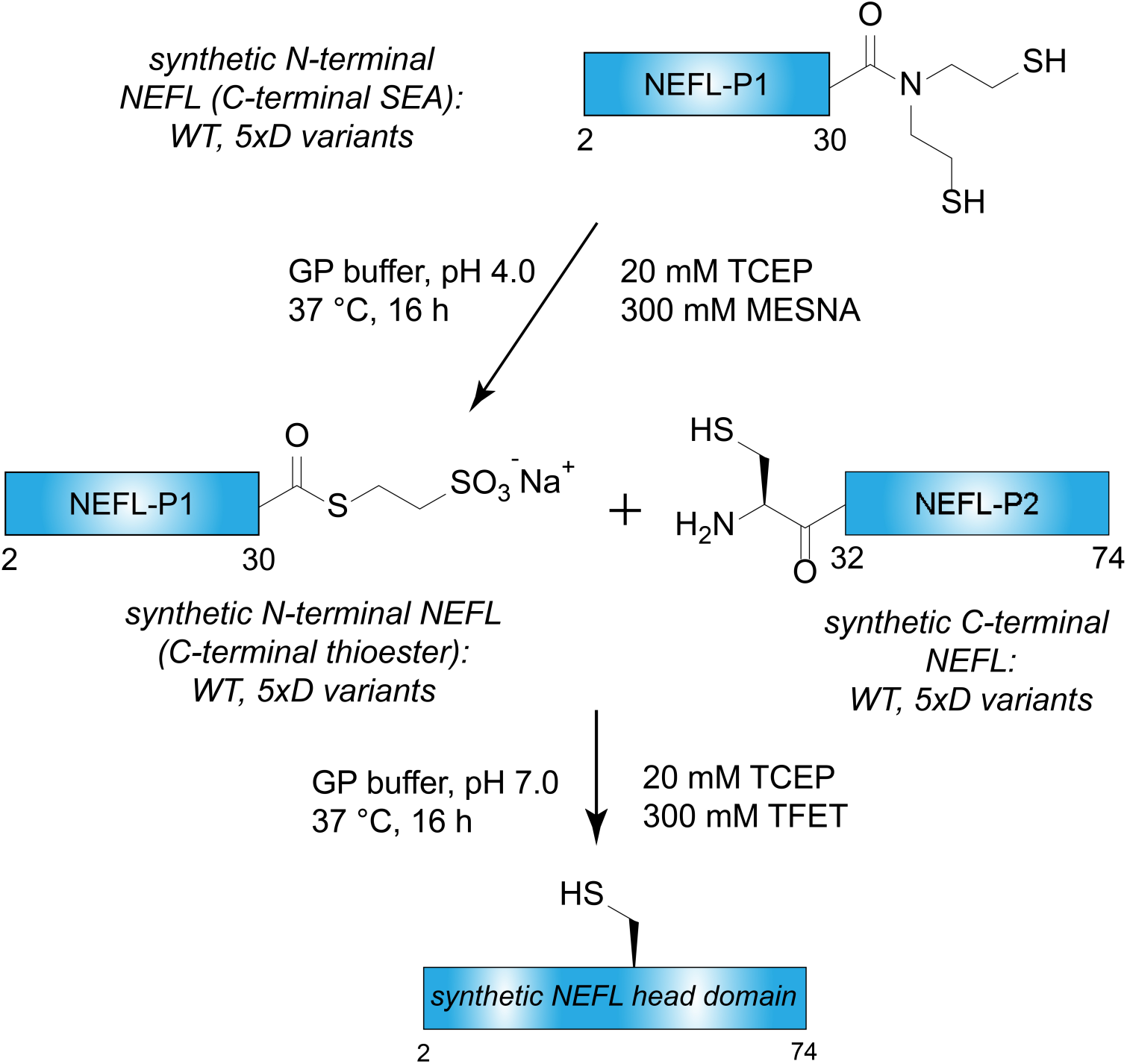
Synthetic strategy to assemble the NEFL LCD. The synthetic scheme used to assemble the NEFL ChIM library. Note: P1 = piece 1; P2 = piece 2.

Previous studies have employed ssNMR to show that, when refolded into physiological buffer conditions, the isolated NEFL head domain adopts a structure identical to that observed in the context of intermediate filaments comprising full-length NEFL protein (8). Accordingly, monitoring head domain self-association provides a robust and experimentally tractable approach to investigate function while circumventing the more complex and material-intensive protocols required for full neurofilament assembly and analysis. In an analogous and straightforward assay for our ChIM scanning campaign, self-association of the isolated head domain can be monitored by solution turbidity at A_395_ (Fig. 4F). We found that N-terminal 5xD constructs, particularly 5xD_22-30_ and 5xD_33-41_ strongly abrogate self-association, with 9.1-fold and 20-fold decreases in turbidity relative to wild-type, respectively. Self-association of 5xD NEFL constructs after 5xD_33-41_, namely 5xD_43-51_, 5xD_53-63_, and 5xD_63-71_, sharply and progressively increases to reach wild-type levels with 2.7-fold, 1.6-fold, and 1-fold relative decreases in turbidity, respectively. Additionally, N-terminal inversions lack smearing due to higher-order oligomer formation characteristic of wild-type head domain when analyzed by SDS-PAGE (Fig. 4D). Thus, ChIM identifies a short, linear motif centered by residues 22-41 within the N-terminus of the NEFL head domain as a region under geometric constraint during self-association.

Prior ssNMR and cryo-EM studies support a model wherein IF head domains engage in labile β-interaction during intermediate filament assembly. CMT-associated NEFL head domain mutation sites (Pro8, Pro22) further implicate the N-terminal portion of the head domain as a site of backbone-mediated intermolecular interactions. Our data complement these studies and suggest that key enantioselective intermolecular interactions are present from residues 22-41 in the NEFL head domain.

## Discussion

LCD self-association activity is a biological phenomenon that regulates cellular signaling processes throughout the cell in both physiological and disease states. To achieve an understanding of LCD biochemistry comparable to that of globular proteins, the chemical properties that govern LCD self-association and interaction specificity must be established with equivalent mechanistic detail and rigor. Here, we apply synthetic protein chemistry to understand how LCDs exploit fundamental principles of the polypeptide backbone to mediate self-association and interaction specificity.

We found that inversion of C^α^ stereochemistry produces significant position-dependent effects on LCD self-association. While many models have been put forth describing how multivalency, bulk amino acid content, and other compositional features underlie LCD function, our data strongly support the existence of labile structural elements that are essential for proper self-association. Consistently, we and others have previously proposed labile cross-β interactions or β-bridging as a common interaction mechanism underlying LCD oligomerization (4, 9, 26–29). We anticipate that this will be a broadly applicable mechanism and emphasize that it is not mutually exclusive with the aforementioned models. Additionally, our data adds to and is consistent with observations of self-association “hotspots” within LCDs of low sequence complexity (9, 26, 30).

We have also demonstrated how synthetic protein chemistry can be leveraged to probe the chemical basis of LCD-LCD interactions with a high degree of positional resolution, which remains challenging when limited to traditional biochemical approaches. We envision that further application of ChIM and synthetic mutagenesis methods will unearth additional LCD sequence features that underlie functional self-association, providing insight into diseases driven by aberrant LCD self-association.

We note that the self-associating segment of the EMD LCD consists of an extended serine homopolymer sequence flanked with aromatic residues. EMD has been proposed to link the nuclear and cytoplasmic cytoskeleton in a phosphorylation- and cell cycle-dependent manner to provide reversible mechanical stability to nuclei (31–33). As such, a comparison may be drawn between the EMD LCD and *Bombyx mori* (mulberry silkworm) sericin, a serine-rich protein “glue” that binds together and waterproofs fibroin filaments in the silkworm cocoon (34). Notably, sericins are comprised of poly-serine stretches interspersed with aromatic and hydrophobic residues and form extensive β-structure networks when reconstituted.

### Limitations of the Study

Here we have demonstrated that synthetic protein mutagenesis offers unique advantages for probing chemical features of the polypeptide backbone that are inaccessible to traditional genetically encoded mutagenesis strategies. However, our approach presents inherent limitations. Compared to genetically encoded mutagenesis, synthetic approaches are material and labor intensive and not as readily compatible with deep scanning methodologies. However, we did find that the ChIM scanning windows chosen for NEFL and EMD were sufficient to identify positional-dependent effects of side chain stereochemical inversion.

Another challenge associated with synthetic protein chemistry is the requirement for unique synthetic pathways for each protein of interest. This includes determination of suitable ligation sites, optimization of synthetic fragment preparation, and potentially optimization of recombinant fragment preparation. These variables are empirically derived and can be time-consuming.

Our study was conducted using reconstituted protein assays because robust ChIM in cellular environments is currently not technically feasible. We addressed this limitation by selecting LCDs with well-established and robust physiological assays: (i) the NEFL head domain’s ability to form structures mirroring those in reconstituted intermediate filaments, and (ii) the established self-immunoprecipitation assays for EMD. For future studies applying this approach to novel proteins, a reliable reconstitution-based assay is a prerequisite. Despite these challenges, the insights gained from ChIM elucidate enantiomerically selective hotspots that enable LCD:LCD interaction specificity.

Lastly, our ChIM analyses of EMD and NEFL each revealed a single, structure-dependent self-association hotspot whose activity is highly sensitive to C^α^ stereochemistry perturbation. While this validates the utility of ChIM to identify geometrically constrained LCD elements, it is important to note that not all LCDs are expected to rely on a single privileged structural motif. LCDs may instead contain multiple, spatially distributed hotspots, each capable of forming chirality-dependent transient structures that collectively contribute to self-association. Additionally, other LCDs may self-associate without forming discrete structure-dependent elements. These concepts should be further explored in future studies of structure-dependent self-association elements in diverse LCD families.

## Materials and Methods

The ChIM and proline insertion mutagenesis variant libraries for EMD and NEFL were generated using a combination of Fmoc solid-phase peptide synthesis, expressed protein ligation, native chemical ligation, and recombinant protein expression in *E. coli*. Recombinant LCD fragments and variants were purified by affinity chromatography followed by reversed-phase HPLC and verified by ESI–LC/MS. Semi-synthetic and fully synthetic LCD proteins were purified by reversed-phase HPLC and verified by ESI–LC/MS. LCD self-association was assessed by turbidity assays following rapid dilution from strong denaturant into physiological buffer conditions. GST-pulldown-based protein–protein interaction assays were analyzed via immunoblotting and densitometric quantification. Detailed materials and methods descriptions are provided in the Supporting Information.

## Acknowledgements

G.L. is funded by the National Institute of General Medical Science (1R35GM147140), the National Cancer Institute (1UM1CA294119), and the Welch Foundation (I-2039-20230405).

## Notes

### Competing Interest Statement

The authors have declared no competing interest.

