## Supporting Information for "Chiral inversion mutagenesis identifies geometrically constrained residues within self-associating low-complexity domains"

**This PDF file includes:**

Supporting Information Methods

Fig. S1-S3

Uncropped Blots

Table S1

### Supporting Information Methods

#### Peptide synthesis

Peptides were synthesized on 0.1 mmol scale on low-loading (0.2 g/mmol) Rink amide PEG/polystyrene ProTide resin (CEM) using automated microwave-assisted Fmoc solid-phase peptide synthesis (SPPS) on a CEM Liberty Blue 2.0 peptide synthesizer (Matthews, NC) with N,N dimethylformamide (DMF, Oakwood Chemical) as solvent. For coupling reactions, 5 molar equivalents (eq) of 9-fluorenylmethyloxycarbonyl (Fmoc)-protected amino acids (Oakwood Chemical) were activated with 5 eq N,N-diisopropylcarbodiimide (DIC, Oakwood Chemical) and 5 eq Oxyma (Oakwood Chemical) and heated to 90 °C for 2 min with $N_2$ bubbling. Fmoc deprotection was carried out with 20% piperidine (Sigma-Aldrich) in DMF supplemented with 0.1 M 1-hydroxybenzotriazole hydrate (HOBt, Oakwood Chemical) at 90 °C for 1 min with $N_2$ bubbling. After synthesis and final Fmoc deprotection, peptides were cleaved from resin with cleavage cocktail of 2.5% $H_2O$, 2.5% triisopropyl silane (TIPS, Sigma), 2.5% ethanedithiol (EDT, Sigma), and 92.5% trifluoroacetic acid (TFA, Oakwood Chemical) for 2 h. Excess TFA was removed by evaporation under atmospheric pressure. Crude peptide cleavage solutions were precipitated with 10 volumes ice-cold diethyl ether (Sigma), centrifuged at 4,000 RCF for 10 min at 4 °C, and dried under atmospheric pressure. The dried peptide pellet was solubilized first by 0.5 mL of neat TFA, followed by addition of aqueous 0.1% TFA and excess guanidinium HCl powder (Fisher Scientific). Solubilized crude peptides were filtered with 0.45-micron syringe filter (Whatman) and injected on a preparative-scale C4 reversed-phase HPLC (RP-HPLC; Agilent) and separated with a 0-70% Solvent B gradient (Solvent A: 0.1% TFA in $H_2O$; Solvent B: 90% acetonitrile in $H_2O$, 0.1% TFA). Fractions were characterized by ESI-LC/MS. Fractions containing pure product (>95%) were pooled and lyophilized (Labconco).

To synthesize NEFL P1 thioester peptides, a modified synthesis procedure was conducted. Bis(2-sulfanylethyl)amido (SEA) polystyrene resin was produced using literature protocols.[1] The first (C-terminal) residue of the NEFL P1 peptide (Fmoc-Arg(Pbf)) was manually coupled to SEA-functionalized polystyrene resin (0.1 mmol/g loading) using 20 eq HATU (Oakwood Chemical), 60 eq N,N-diisopropylethylamine (DIPEA, Sigma), and 20 eq Fmoc-Arg(Pbf) in DMF for 30 min at 21°C. This loading protocol was repeated. After second coupling, resin was washed with DMF and acetyl-capped with excess acetic anhydride and DIPEA in 5 mL DMF for 10 min. The remainder of synthesis was conducted using automated microwave SPPS as described above. After synthesis, peptides were cleaved from resin with a cleavage cocktail containing 2.5% $H_2O$, 2.5% TIPS, and 95% TFA (without EDT) for 2 h. After cleavage and precipitation, the C-terminal SEA moiety was converted to a sodium-2-mercaptoethanesulfonate (MesNa) thioester by incubating the dried crude peptide pellet in aqueous buffer containing 6 M guanidium hydrochloride, 100 mM sodium phosphate pH 4.0 ("GP buffer"), 20 mM tris(2-carboxyethyl)phosphine (TCEP, GoldBio) and 300 mM MesNa (Sigma) 12 h at 37 °C. MesNa thioester peptides were purified by preparative C4 RP-HPLC as described above.

**Molecular cloning**

Standard molecular cloning techniques were employed to generate recombinant expression plasmids. EMD LCD was subcloned from human EMD cDNA purchased from Ultimate™ ORF Lite human cDNA collection (Life Technologies) using Gibson assembly into pET30 vectors to generate 6xHis-SUMO-EMD_LCD and 6xHis-SUMO-EMD_LCD-GyrA-6xHis constructs or into the pGex vector to create the GST-EMD_LCD construct. Proline insertion mutagenesis and addition of hemagglutinin (HA) tag was performed with inverse PCR site directed mutagenesis. Primers for cloning were synthesized by Integrated DNA Technologies (IDT).

**Protein purification**

Recombinant proteins were expressed in BL21 Rosetta (DE3) *E. coli* cells (Novagen) using standard methods. Primary cultures in Luria broth (LB) were inoculated with *E. coli* transformed with expression plasmids and incubated for 16 h at 37 °C. Expression cultures (1 L) were inoculated with 2 mL each of primary culture and incubated at 37 °C until reaching $OD_{600}$ of 0.5. The incubation temperature was then lowered to 16 °C and cells were induced with 0.1 mg/mL IPTG (GoldBio) for 16 h. Cell pellets were collected with centrifugation at 4,000 RCF.

For recombinant full-length EMD LCD constructs (residues 46-222, 6xHis-SUMO fusion), cell pellets from 2 L expression culture were lysed via sonication (75% amplitude, 5 sec on, 15 sec off, 3 min total) in denaturing lysis buffer (50 mM Tris-HCl pH 7.2, 100 mM NaCl, 20 mM imidazole, 5 mM beta-mercaptoethanol (BME), 1 mM phenylmethylsulfonyl fluoride (PMSF), 6 M urea). Lysate was clarified with centrifugation and applied to equilibrated Ni-NTA resin (Qiagen) at 4 °C for 1 h. Resin was washed with 5 CV of denaturing lysis buffer. Resin was then washed with an additional 2 CV of lysis buffer containing 2 M urea and eluted with 10 mL of 2 M urea lysis buffer supplemented with 300 mM imidazole and 1 mM TCEP. Ulp1 protease was added to eluent and incubated at 4 °C for 1 h. After Ulp1 proteolysis, the solution was syringe-filtered with a 0.45 micron filter and purified with preparative C4 RP-HPLC (0-70% Solvent B gradient over 30 min). Fractions containing pure product (>95% purity) were characterized by ESI-LC/MS, pooled, and lyophilized. The GST-EMD LCD protein was expressed as described above with lysis conducted in lysis buffer lacking denaturant.

To produce EMD-LCD P1 thioester, an EMD LCD fragment (residues 46-188) was expressed with an N-terminal 6xHis-tagged Sumo fusion and a C-terminal 6xHis-tagged Gyrase A intein fusion. Cell pellets were lysed in non-denaturing lysis buffer (50 mM Tris-HCl pH 7.2, 500 mM NaCl, 20 mM imidazole, 1 mM PMSF, no reducing agent) via sonication. Lysate was clarified with centrifugation and applied to Ni-NTA resin. Resin was washed with 5 CV of lysis buffer and eluted with 20 mL lysis buffer supplemented with 300 mM imidazole. Eluent was then dialyzed for 12 h at 4 °C using 3.5 kDa cutoff cellulose tubing (Repligen) against lysis buffer without imidazole. To initiate Gyrase A thiolysis, 200mM MesNa and 10 mM pH-neutralized TCEP were added to the dialyzed eluent solution and pH adjusted to 7.0. The eluent solution was then incubated for 16 h at 21 °C. The following day, Ulp1 protease was added to the solution and incubated at 21°

C for an additional 1 h. The thiolysis solution was then syringe-filtered and purified with preparative C4 RP-HPLC (0-70% Solvent B gradient over 30 min). Fractions containing pure product were characterized by ESI-LC/MS, pooled, and lyophilized.

**Protein semi-synthesis**

To prepare semi-synthetic EMD LCD, lyophilized P1 recombinant thioester and P2 synthetic N-terminal thiol peptide fragment were solubilized in $N_2$-degassed GP ligation buffer (6 M guanidium-HCl, 100 mM sodium phosphate pH 7.0) to concentrations of 0.5 mM and 2 mM, respectively. The solution was supplemented with an additional 10 mM TCEP and 300 mM 2,2,2-trifluoroethanethiol (TFET) and the pH re-adjusted to 7.0. Ligation solutions were sealed with parafilm and incubated at 37° C for 16 h. Following the incubation, ligation solutions were supplemented with an additional 2 mM TCEP and purified via semi-preparative C4 RP-HPLC using a 20-40% Solvent B gradient over 30 min. Peaks were collected manually according to absorbance at 214nm. Fractions containing pure ligation product were lyophilized.

To prepare synthetic NEFL head domain, lyophilized P1 and P2 peptides were solubilized in GP ligation buffer to 0.5 mM. Reactions were immediately supplemented with 10 mM TCEP and 300 mM TFET and the pH adjusted to 7.0 as above. Purification procedures were conducted as above via C4 RP-HPLC with 0-50% Solvent B gradient over 30 min.

**Turbidity assays**

Lyophilized EMD-LCD variants were resuspended to 1.2 mM (3xPro) or 1 mM (3xD) in 25 mM Tris-HCl buffer pH (7.0) with 6 M urea. Resuspended EMD-LCD solutions were then diluted 1:20 into non-denaturing buffer (50 mM Tris-HCl, 50 mM NaCl, 1 mM DTT), mixed with pipetting and placed in a clear half-volume 96-well plate. Turbidity of the solution was measured immediately at 395 nm wavelength using a Cytation 5 microplate reader (Biotek).

For NEFL turbidity assays, lyophilized NEFL head domains were resuspended to 2 mM in 25 mM Tris-HCl buffer (pH 7.0) containing 6 M urea. NEFL head domain solutions were then diluted 1:20 into high-salt non-denaturing buffer (50 mM Tris-HCl, 300 mM NaCl, 1 mM DTT), mixed with pipetting and placed in a clear half-volume 96-well plate. Turbidity of the solution was measured immediately at 395 nm wavelength using a Cytation 5 microplate reader (Biotek).

**EMD GST pulldown**

Clarified lysate from 1 L GST-EMD LCD expression culture was incubated with GSH agarose resin (Qiagen) for 1 hour at 4 °C. Resin was then washed extensively with 10 eq of non-denaturing lysis buffer. Resin was then washed with EMD binding buffer adapted from prior literature (300 mM NaCl, 50 mM Tris-HCl pH 7.3, 0.1% Triton, 1 mM DTT).[2] GST-EMD LCD-loaded resin was aliquoted to 25 uL of packed resin per pulldown.

Lyophilized HA-tagged EMD LCD variants were resuspended in 25 mM Tris-HCl, 6 M urea buffer to a concentration of 10 uM. HA-EMD LCD solutions were further diluted 1:200 in non-denaturing EMD binding buffer to yield a final concentration of 50 nM. HA-tagged EMD LCD was applied to the immobilized GST-EMD LCD and incubated at 4 °C for 1 h. Bound resin was washed with 4 CV of EMD binding buffer and elution was carried out via incubation in EMD binding buffer supplemented with 10 mM GSH for 10 min at 4 °C. Input (1:5 dilution) and eluent solutions were then loaded on an SDS-PAGE gel and transferred to a PVDF membrane for Western blot. Membranes were probed by anti-HA primary at a 1:2000 dilution (Cell Signaling Technologies, HA-Tag (C29F4) Rabbit mAb #3724), washed with PBST and ultimately imaged via a fluorophore-conjugated secondary antibody. Membranes were imaged on a BioRad ChemiDoc imaging system. Fraction pulldown was assessed by comparing band density as quantified with ImageJ densitometry.

**Peptide sequences**

**lowercase :** D-amino acids

**EMD P2 peptides:**
EMD WT:
CFMSSSSSSSSWLTRRAIRPENRAPGAGLGQER

EMD 3xD 191-195:
CF**m**S**s**S**s**SSSSWLTRRAIRPENRAPGAGLGQER

EMD 3xD 195-199
CFMSSS**s**S**s**S**s**WLTRRAIRPENRAPGAGLGQER

EMD 3xD 199-203
CFMSSSSSSS**s**W**l**T**r**RAIRPENRAPGAGLGQER

EMD 3xD 203-207
CFMSSSSSSSSWLT**r**R**a**I**r**PENRAPGAGLGQER

EMD 3xD 207-211
CFMSSSSSSSSWLTRRAI**r**P**e**N**r**APGAGLGQER

EMD 3xD 211-215
CFMSSSSSSSSWLTRRAIRPEN**r**A**p**G**a**GLGQER

**NEFL P1 peptides:**
NEFL WT P1
SSFSYEPYYSTSYKRRYVETPRVHISSVR-MesNa

NEFL 5xD 2-10
**s**S**f**S**y**E**p**Y**y**STSYKRRYVETPRVHISSVR-MesNa

NEFL 5xD 12-20
SSFSYEPYYS**t**S**y**K**r**R**y**V**e**TPRVHISSVR-MesNa

NEFL 5xD 22-30
SSFSYEPYYSTSYKRRYVET**p**R**v**H**i**S**s**V**r**-MesNa

**NEFL P2 peptides:**
NEFL WT P2
CGYSTARSAYSSYSAPVSSSLSVRRSYSSSSGSLMPSLENLELS

NEFL 5xD 33-41
CG**y**S**t**A**r**S**a**Y**s**SYSAPVSSSLSVRRSYSSSSGSLMPSLENLELS

NEFL 5xD 43-51
CGYSTARSAYSS**y**S**a**P**v**S**s**S**l**SVRRSYSSSSGSLMPSLENLELS

NEFL 5xD 53-61
CGYSTARSAYSSYSAPVSSSLS**v**R**r**S**y**S**s**S**s**GSLMPSLENLELS

NEFL 5xD 63-71
CGYSTARSAYSSYSAPVSSSLSVRRSYSSSSG**s**L**m**P**s**L**e**N**l**ELS

**Protein sequences:**
**EMD 3xPro proteins:**
EMD LCD WT
(YPYDVPDYASG)RRLSPPSSSAASSYSFSDLNSTRGDADMYDLPKKEDALLYQSKGYNDDYY
EESYFTTRTYGEPESAGPSRAVRQSVTSFPDADAFHHQVHDDDLLSSSEEECKDRERPMYGR
DSAYQSITHYRPVSASRSSLDLSYYPTSSSTSFMSSSSSSSSWLTRRAIRPENRAPGAGLGQD
RQ

EMD LCD 3xPro 143-146
RRLSPPSSSAASSYSFSDLNSTRGDADMYDLPKKEDALLYQSKGYNDDYYEESYFTTRTYGEP
ESAGPSRAVRQSVTSFPDADAFHHQVHDDDLLSSS**P**E**P**E**P**ECKDRERPMYGRDSAYQSITHY
RPVSASRSSLDLSYYPTSSSTSFMSSSSSSSSWLTRRAIRPENRAPGAGLGQDRQ

EMD LCD 3xPro 153-156
RRLSPPSSSAASSYSFSDLNSTRGDADMYDLPKKEDALLYQSKGYNDDYYEESYFTTRTYGEP
ESAGPSRAVRQSVTSFPDADAFHHQVHDDDLLSSSEEECKDRERP**P**M**P**Y**P**GRDSAYQSITHY
RPVSASRSSLDLSYYPTSSSTSFMSSSSSSSSWLTRRAIRPENRAPGAGLGQDRQ

EMD LCD 3xPro 163-166
RRLSPPSSSAASSYSFSDLNSTRGDADMYDLPKKEDALLYQSKGYNDDYYEESYFTTRTYGEP
ESAGPSRAVRQSVTSFPDADAFHHQVHDDDLLSSSEEECKDRERPMYGRDSAYQS**P**I**P**T**P**HY
RPVSASRSSLDLSYYPTSSSTSFMSSSSSSSSWLTRRAIRPENRAPGAGLGQDRQ

EMD LCD 3xPro 173-176
RRLSPPSSSAASSYSFSDLNSTRGDADMYDLPKKEDALLYQSKGYNDDYYEESYFTTRTYGEP
ESAGPSRAVRQSVTSFPDADAFHHQVHDDDLLSSSEEECKDRERPMYGRDSAYQSITHYRPV
SAS**P**R**P**S**P**SLDLSYYPTSSSTSFMSSSSSSSSWLTRRAIRPENRAPGAGLGQDRQ

(HA)EMD LCD 3xPro 178-181
(YPYDVPDYASG)RRLSPPSSSAASSYSFSDLNSTRGDADMYDLPKKEDALLYQSKGYNDDYY
EESYFTTRTYGEPESAGPSRAVRQSVTSFPDADAFHHQVHDDDLLSSSEEECKDRERPMYGR
DSAYQSITHYRPVSASRSSLD**P**L**P**S**P**YYPTSSSTSFMSSSSSSSSWLTRRAIRPENRAPGAGL
GQDRQ

(HA)EMD LCD 3xPro 183-186
(YPYDVPDYASG)RRLSPPSSSAASSYSFSDLNSTRGDADMYDLPKKEDALLYQSKGYNDDYY
EESYFTTRTYGEPESAGPSRAVRQSVTSFPDADAFHHQVHDDDLLSSSEEECKDRERPMYGR
DSAYQSITHYRPVSASRSSLDLSYYP**P**T**P**S**P**SSTSFMSSSSSSSSWLTRRAIRPENRAPGAGL
GQDRQ

(HA)EMD LCD 3xPro 188-191

(YPYDVPDYASG)RRLSPPSSSAASSYSFSDLNSTRGDADMYDLPKKEDALLYQSKGYNDDYY
EESYFTTRTYGEPESAGPSRAVRQSVTSFPDADAFHHQVHDDDLLSSSEEECKDRERPMYGR
DSAYQSITHYRPVSASRSSLDLSYYPTSSST**P**S**P**F**P**MSSSSSSSSWLTRRAIRPENRAPGAGL
GQDRQ

(HA)EMD LCD 3xPro 193-196
(YPYDVPDYASG)RRLSPPSSSAASSYSFSDLNSTRGDADMYDLPKKEDALLYQSKGYNDDYY
EESYFTTRTYGEPESAGPSRAVRQSVTSFPDADAFHHQVHDDDLLSSSEEECKDRERPMYGR
DSAYQSITHYRPVSASRSSLDLSYYPTSSSTSFMSS**P**S**P**S**P**SSSSWLTRRAIRPENRAPGAGL
GQDRQ

(HA)EMD LCD 3xPro 198-201
(YPYDVPDYASG)RRLSPPSSSAASSYSFSDLNSTRGDADMYDLPKKEDALLYQSKGYNDDYY
EESYFTTRTYGEPESAGPSRAVRQSVTSFPDADAFHHQVHDDDLLSSSEEECKDRERPMYGR
DSAYQSITHYRPVSASRSSLDLSYYPTSSSTSFMSSSSSSS**P**S**P**W**P**LTRRAIRPENRAPGAGL
GQDRQ

(HA)EMD LCD 3xPro 203-206
(YPYDVPDYASG)RRLSPPSSSAASSYSFSDLNSTRGDADMYDLPKKEDALLYQSKGYNDDYY
EESYFTTRTYGEPESAGPSRAVRQSVTSFPDADAFHHQVHDDDLLSSSEEECKDRERPMYGR
DSAYQSITHYRPVSASRSSLDLSYYPTSSSTSFMSSSSSSSSWLTR**P**R**P**A**P**IRPENRAPGAGL
GQDRQ

(HA)EMD LCD 3xPro 208-211
(YPYDVPDYASG)RRLSPPSSSAASSYSFSDLNSTRGDADMYDLPKKEDALLYQSKGYNDDYY
EESYFTTRTYGEPESAGPSRAVRQSVTSFPDADAFHHQVHDDDLLSSSEEECKDRERPMYGR
DSAYQSITHYRPVSASRSSLDLSYYPTSSSTSFMSSSSSSSSWLTRRAIRP**P**E**P**N**P**RAPGAGL
GQDRQ

(HA)EMD LCD 3xPro 213-216
(YPYDVPDYASG)RRLSPPSSSAASSYSFSDLNSTRGDADMYDLPKKEDALLYQSKGYNDDYY
EESYFTTRTYGEPESAGPSRAVRQSVTSFPDADAFHHQVHDDDLLSSSEEECKDRERPMYGR
DSAYQSITHYRPVSASRSSLDLSYYPTSSSTSFMSSSSSSSSWLTRRAIRPENRAP**P**G**P**A**P**GL
GQDRQ

(HA)EMD P1 fragment
(YPYDVPDYASG)RRLSPPSSSAASSYSFSDLNSTRGDADMYDLPKKEDALLYQSKGYNDDYY
EESYFTTRTYGEPESAGPSRAVRQSVTSFPDADAFHHQVHDDDLLSSSEEECKDRERPMYGR
DSAYQSITHYRPVSASRSSLDLSYYPTSSST-MesNa

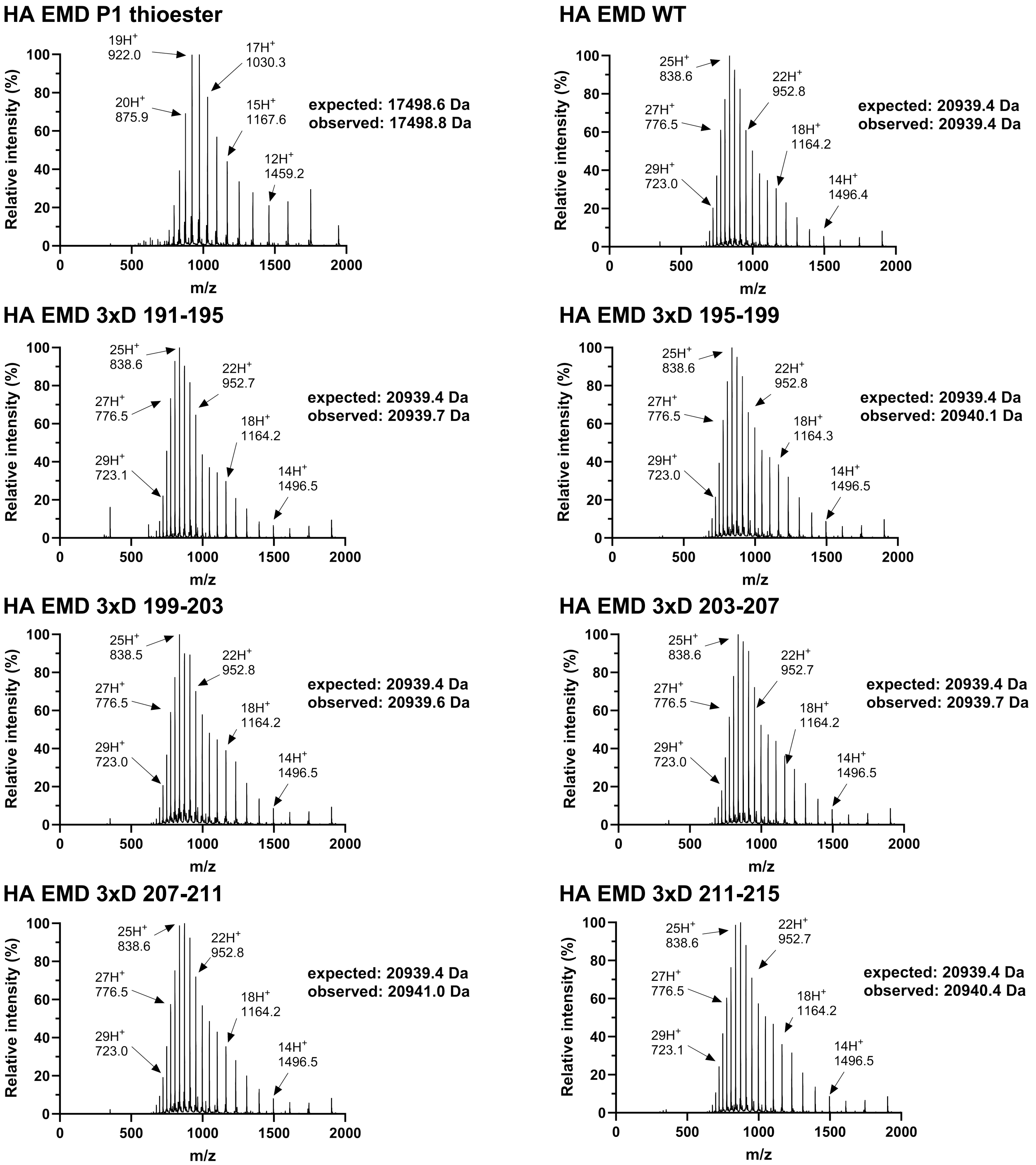


**Fig. S1. ESI-LC/MS characterization of HA-tagged semi-synthetic EMD LCD.**
Indicated spectra are derived from whole-peak integration of ESI-LC/MS trace. For HA P1 thioester, an additional species with mass of 16145.6 Da corresponding to thioester hydrolysis can be observed (10%).

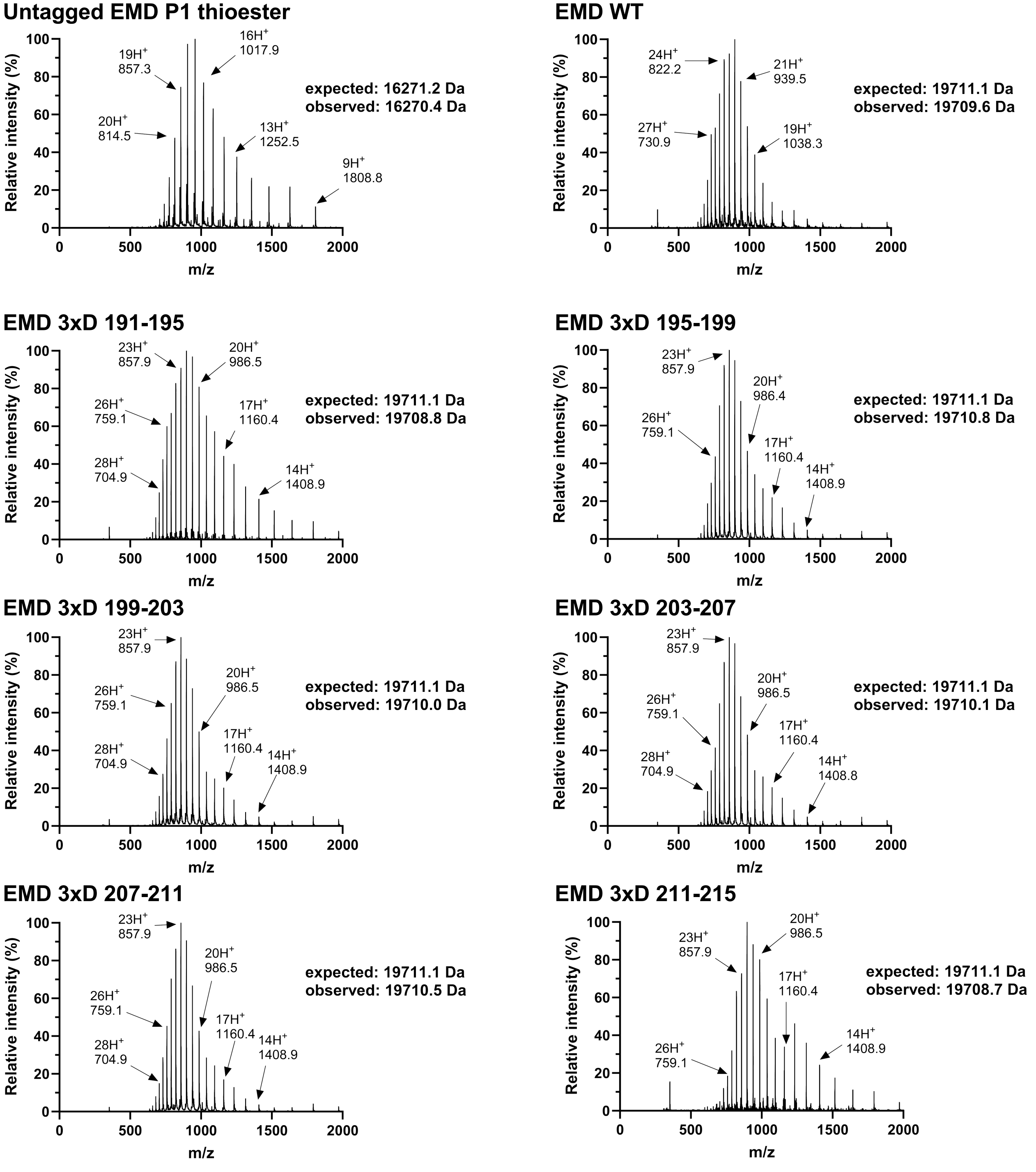


**Fig. S2. ESI-LC/MS characterization of untagged semi-synthetic EMD LCD**
Indicated spectra are derived from whole-peak integration of ESI-LC/MS trace. For untagged P1 thioester, an additional species with mass of 16145.6 Da corresponding to thioester hydrolysis can be observed (15%).

**NEFL WT**

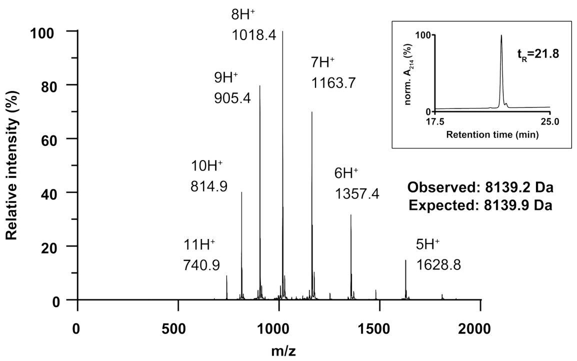


**NEFL 5xD 2-10**

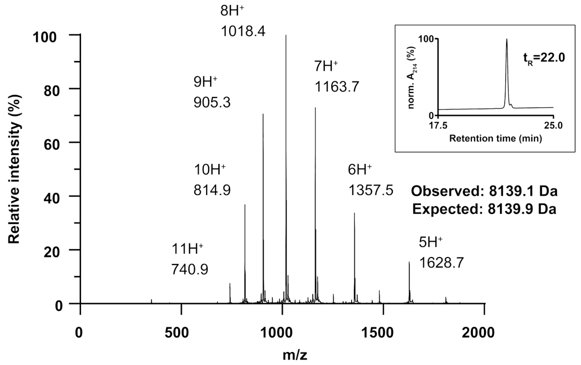


**NEFL 5xD 12-20**

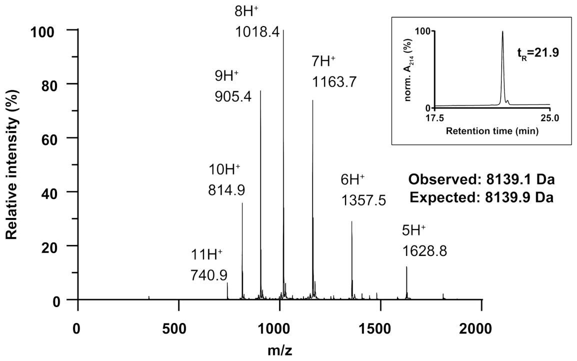


**NEFL 5xD 22-30**

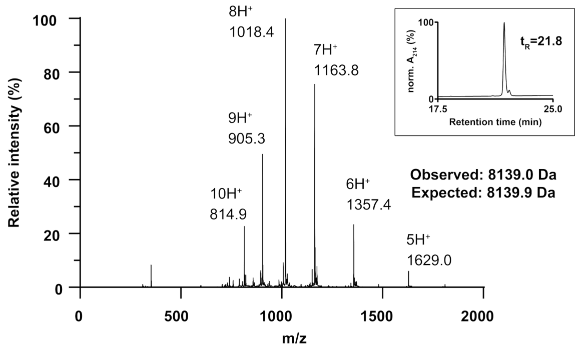


**NEFL 5xD 43-51**

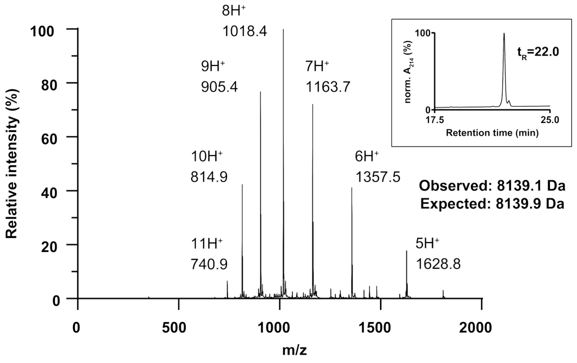


**NEFL 5xD 53-61**

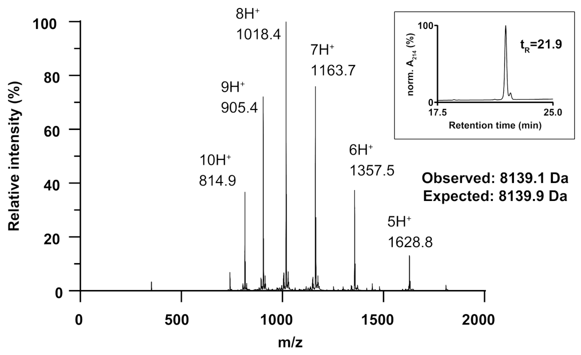


**NEFL 5xD 63-71**

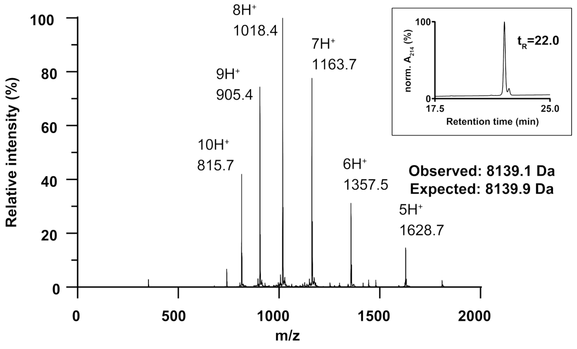


**Fig. S3: Synthetic NEFL head domain characterization**

ESI-LC/MS and C4 analytical RP-HPLC (inset) characterization of indicated synthetic head domains. For all samples, a species with +16 Da mass shift can also be observed between ~15-20% of target mass abundance, likely due to methionine oxidation. This adduct was also observed with purified recombinant NEFL head domain.

**Uncropped Blots**

Blots used for Fig. 1D quantification:

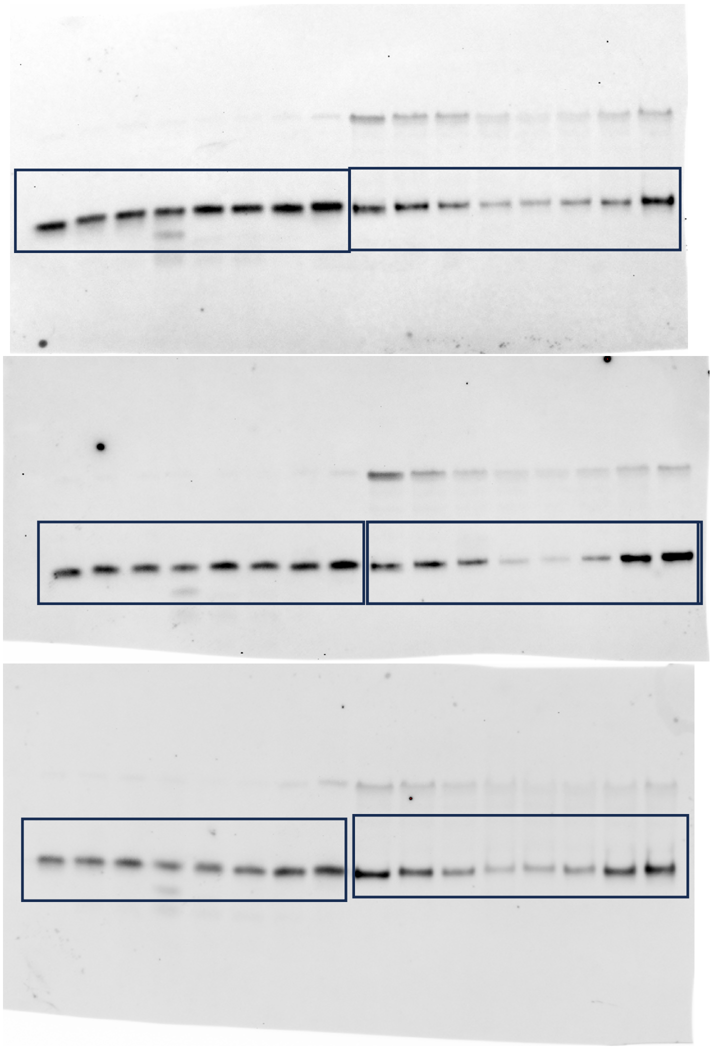

Blots used for Fig. 2E:

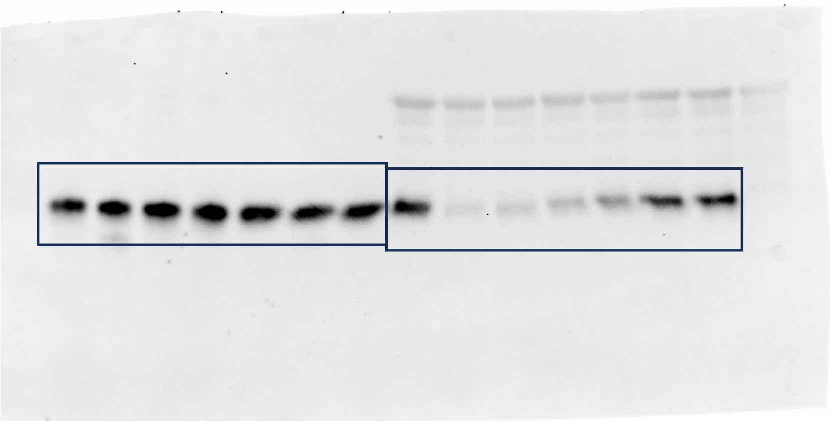

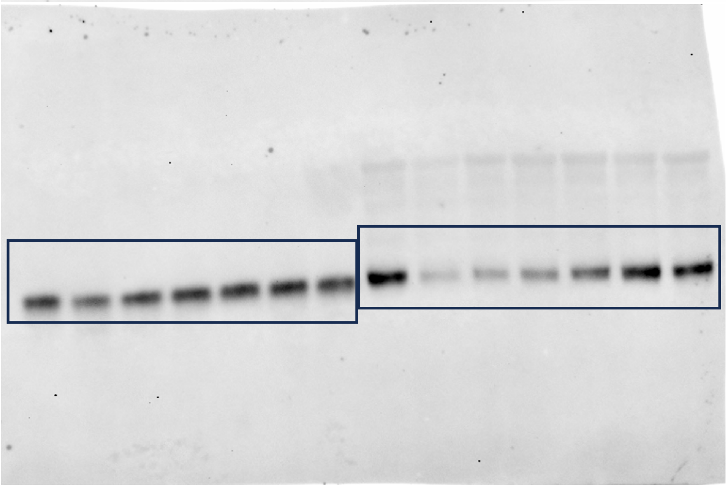

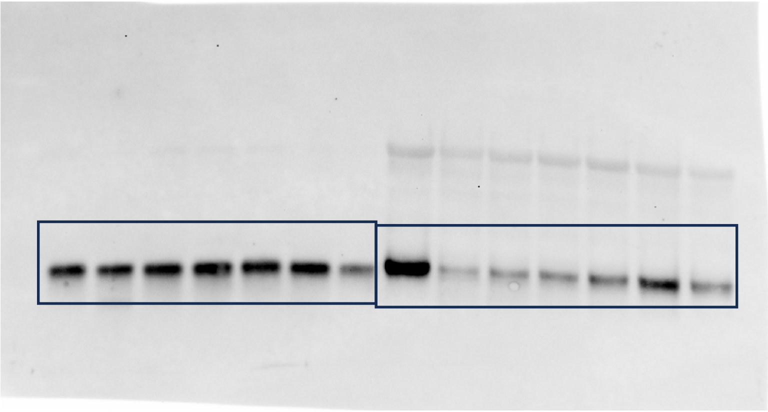

**Table S1. Key resources table**

| REAGENT OR RESOURCE | SOURCE | IDENTIFIER |
|---|---|---|
| **Antibodies** | | |
| Anti-HA monoclonal | Cell Signaling Technologies | C29F4 |
| Goat anti-Mouse IgG (H+L) Cross-Adsorbed Secondary Antibody, Alexa Fluor™ 488 | Thermo Fisher Scientific | A-11001 |
| Goat anti-Rabbit IgG (H+L) Cross-Adsorbed Secondary Antibody, Alexa Fluor™ 647 | Thermo Fisher Scientific | A-21244 |
| **Bacterial cell lines** | | |
| Mach1 (*E. coli*) | Thermo Fisher Scientific | C862003 |
| Rosetta2 (*E. coli*) | MilliporeSigma | 71402 |
| **Recombinant DNA, cloning reagents** | | |
| Phusion High-Fidelity DNA Polymerase | NEB | M0530L |
| Kapa HiFi HotStart ReadyMix | Roche | 9420398001 |
| Gibson Assembly Master Mix | NEB | E2611L |
| DpnI nuclease | NEB | R0176 |
| human EMD cDNA | Horizon Discovery | MHS6278-202828259 |
| pET30 HPF1-His-SUMO-Flag | Addgene | 111577 |
| pET30 6xHis-SUMO-EMD LCD | This study | N/A |
| pET30 6xHis-SUMO-EMD LCD 3xPro 143-146 | This study | N/A |
| pET30 6xHis-SUMO-EMD LCD 3xPro 153-156 | This study | N/A |
| pET30 6xHis-SUMO-EMD LCD 3xPro 163-166 | This study | N/A |
| pET30 6xHis-SUMO-EMD LCD 3xPro 173-176 | This study | N/A |
| pET30 6xHis-SUMO-EMD LCD 3xPro 178-181 | This study | N/A |
| pET30 6xHis-SUMO-EMD LCD 3xPro 183-186 | This study | N/A |
| pET30 6xHis-SUMO-EMD LCD 3xPro 188-191 | This study | N/A |
| pET30 6xHis-SUMO-EMD LCD 3xPro 193-196 | This study | N/A |
| pET30 6xHis-SUMO-EMD LCD 3xPro 198-201 | This study | N/A |
| pET30 6xHis-SUMO-EMD LCD 3xPro 203-206 | This study | N/A |
| pET30 6xHis-SUMO-EMD LCD 3xPro 208-211 | This study | N/A |
| pET30 6xHis-SUMO-EMD LCD 3xPro 213-216 | This study | N/A |
| pET30 6xHis-SUMO-HA-EMD LCD | This study | N/A |

| | | |
|---|---|---|
| pET30 6xHis-SUMO-HA-EMD LCD 3xPro 178-181 | This study | N/A |
| pET30 6xHis-SUMO-HA-EMD LCD 3xPro 183-186 | This study | N/A |
| pET30 6xHis-SUMO-HA-EMD LCD 3xPro 188-191 | This study | N/A |
| pET30 6xHis-SUMO-HA-EMD LCD 3xPro 193-196 | This study | N/A |
| pET30 6xHis-SUMO-HA-EMD LCD 3xPro 198-201 | This study | N/A |
| pET30 6xHis-SUMO-HA-EMD LCD 3xPro 203-206 | This study | N/A |
| pET30 6xHis-SUMO-HA-EMD LCD 3xPro 208-211 | This study | N/A |
| pET30 6xHis-SUMO-HA-EMD LCD 3xPro 213-216 | This study | N/A |
| pET30 6xHis-SUMO-HA-EMD LCD 3xPro 178-181 | This study | N/A |
| pET30 6xHis-SUMO-HA-EMD LCD 3xPro 183-186 | This study | N/A |
| pET30 6xHis-SUMO-EMD P1-GyrA-6xHis | This study | N/A |
| pET30 6xHis-SUMO-HA-EMD P1-GyrA-6xHis | This study | N/A |
| pGEX GST-EMD LCD | This study | N/A |
| **Software** | | |
| Prism 10.0.0 | GraphPad | https://www.graphpad.com/ |
| Adobe Illustrator | Adobe | https://www.adobe.com/products/illustrator.html |
| BioTek Gen5 | Agilent Technologies, Inc | https://www.agilent.com/en/product/cell-analysis/cell-imaging-microscopy/cell-imaging-microscopy-software |
| Agilent OpenLab ChemStation | Agilent Technologies, Inc | https://www.agilent.com/en/product/software-informatics/analytical-software-suite/chromatography-data-systems/openlab-chemstation |

| | | |
|---|---|---|
| ImageJ | NIH | https://imagej.net/ij/ |
| **Chemicals, synthetic peptides, recombinant proteins** | | |
| Acetonitrile | MilliporeSigma | 34851 |
| N,N-dimethylformamide | Fisher Scientific | D119 |
| Rink Amide ProTide (LL) resin | CEM | R003-B |
| SEA-polystyrene resin | This study, ref 1. | N/A |
| Fmoc-protected amino acids | Oakwood Chemical | N/A |
| Trifluoroacetic acid | Oakwood Chemical | 001271 |
| Formic acid | Fisher Scientific | A117 |
| N,N′-diisopropylcarbodiimide | Oakwood Chemical | M02889 |
| Oxyma | Oakwood Chemical | 043278 |
| Piperidine | MilliporeSigma | 104094 |
| 1-hydroxybenzotriazole hydrate | Oakwood Chemical | M02875 |
| HATU | Oakwood Chemical | 023926 |
| N,N-diisopropylethylamine | MilliporeSigma | O4884 |
| Triisopropylsilane | MilliporeSigma | 233781 |
| Acetic anhydride | Oakwood Chemical | 035908 |
| 1,2-ethanedithiol | MilliporeSigma | 02390 |
| TCEP-HCl | GoldBio | TCEP |
| Glutathione, reduced | Sigma | G6013 |
| DTT | Research Products Intl. | D11000 |
| Imidazole | Sigma | I5513 |
| IPTG | GoldBio | I2481C |
| Luria broth, granulated | Fisher Scientific | BP97235 |
| PMSF | GoldBio | P-470 |
| **Commercial assays, kits, and resins** | | |
| Pierce Glutathione Agarose | ThermoFisher | 16101 |
| HisPUR Ni-NTA Resin | ThermoFisher | 88221 |

### Supporting Information References

(1) Ollivier, N.; Raibaut, L.; Blanpain, A.; Desmet, R.; Dheur, J.; Mhidia, R.; Boll, E.; Drobecq, H.; Pira, S. L.; Melnyk, O. Tidbits for the Synthesis of Bis(2-Sulfanylethyl)Amido (SEA) Polystyrene Resin, SEA Peptides and Peptide Thioesters. *Journal of Peptide Science* **2014**, *20* (2), 92–97. https://doi.org/10.1002/psc.2580.

(2) Berk, J. M.; Simon, D. N.; Jenkins-Houk, C. R.; Westerbeck, J. W.; Grønning-Wang, L. M.; Carlson, C. R.; Wilson, K. L. The Molecular Basis of Emerin-Emerin and Emerin-BAF Interactions. *J Cell Sci* **2014**, *127* (Pt 18), 3956–3969. https://doi.org/10.1242/jcs.148247.
